# Cell Painting for cytotoxicity and mode-of-action analysis in primary human hepatocytes

**DOI:** 10.1101/2025.01.22.634152

**Authors:** Jessica D. Ewald, Katherine L. Titterton, Alex Bäuerle, Alex Beatson, Daniil A. Boiko, Ángel A. Cabrera, Jaime Cheah, Beth A. Cimini, Bram Gorissen, Thouis Jones, Konrad J. Karczewski, David Rouquie, Srijit Seal, Erin Weisbart, Brandon White, Anne E. Carpenter, Shantanu Singh

**Affiliations:** - Imaging Platform, Broad Institute of MIT and Harvard, Cambridge MA, USA; - Axiom Bio, San Francisco CA, USA; - The Center for the Development of Therapeutics, Broad Institute of MIT and Harvard, Cambridge MA, USA; - The Novo Nordisk Foundation Center for Genomic Mechanisms of Disease, Broad Institute of MIT and Harvard, Cambridge MA, USA; - Toxicology Data Science, Bayer SAS Crop Science Division, Valbonne Sophia-Antipolis, France

## Abstract

High-throughput, human-relevant approaches for predicting chemical toxicity are urgently needed for better decision-making in human health. Here, we apply image-based profiling (the Cell Painting assay) and two cytotoxicity assays (metabolic and membrane damage readouts) to primary human hepatocytes after exposure to eight concentrations of 1085 compounds that include pharmaceuticals, pesticides, and industrial chemicals with known liver toxicity-related outcomes. Three computational methods (CellProfiler, a Cell Painting-specific convolutional neural network, and a pretrained vision transformer) were compared to extract morphology features from single cells or entire images. We used these morphology features to predict activity in the measured cytotoxicity assays, as well as in 412 curated ToxCast assays that span cytotoxicity, cell-based, and cell-free categories. We found that the morphological profiles detect compound bioactivity at lower concentrations than standard cytotoxicity assays. In supervised analyses, they predict cytotoxicity and targeted cell-based assay readouts, but not cell-free assay readouts. We also found that the various feature extraction methods performed relatively similarly and that filtering out non-bioactive or cytotoxic concentrations did not boost supervised assay prediction performance for any assay endpoint category, although it did have a large influence on unsupervised cluster analysis. We envision that image-based profiling could serve as a key component of modern safety assessment.

## Introduction

High-throughput approaches for predicting *in vivo* toxicity of compounds in humans are urgently needed. The majority of the >350,000 compounds in commercial use worldwide have never been assessed for adverse impacts on human health ^1^. Even for pharmaceuticals, which undergo extensive preclinical testing, many fail clinical trials due to toxicities not predicted by animal models, draining resources and increasing drug development costs ^2^. Animal-based testing is resource-intensive, raises ethical concerns, and often fails to accurately predict outcomes in humans, making them insufficient for addressing these data gaps. Thus, scientists in government, industry, and academia are working to bring about a new paradigm in which a suite of human-relevant *in silico* and *in vitro* models are used to understand and predict the *in vivo* impacts of chemical exposures at scale ^3,4^.

*In vitro* cellular models are relevant to *in vivo* toxicity when we can link a compound exposure to changes in cells which themselves have solid associations to organism-level toxicity, for example binding to a specific receptor, initiating metabolic reprogramming, or triggering a particular cell death pathway. In the short term, the objective is to identify cellular perturbations that are obviously relevant to classical toxicity outcomes of interest. For example, *in vitro* toxicity testing methods have been developed to assess the skin sensitization ^5^ or genotoxic ^6^ potential of compounds. In the long term, the goal is to define causal relationships between compound-induced perturbations and key impacts across biological scales, from molecules to cells to tissues to organisms, also called “adverse outcome pathways” ^7,8^. If we have high confidence in how a particular perturbation at the cellular level manifests as toxicity at the organism level, then observing this perturbation *in vitro* should be a solid basis for screening chemical toxicity reliably at scale. Achieving this vision requires high-throughput methods that can identify diverse impacts of compound exposure on cell state across a range of concentrations.

Targeted *in vitro* assays are one way of obtaining readouts indicative of specific cell states, however it is logistically difficult to run enough assays in parallel to comprehensively cover biological space. The ToxCast/Tox21 program screened ∼900 targeted assay endpoints for >8900 compounds ^9^; while this is an extremely valuable source of information, it has been a nearly 20-year effort from initial experimental design to the public release of processed data ^10,11^. More recent strategic approaches call for the initial collection of high-throughput, high-dimensional omics data that capture broad swathes of biological space in a single assay ^4^. The Cell Painting assay is a promising approach for acquiring high-throughput omics data from cells that might be predictive of specific toxicity-related outcomes ^12^. It is an image-based profiling assay using six fluorescent dyes to label eight different cell components, and it measures thousands of features, including intensity, shape, and texture of the various stains in various regions of the cell ^13^. It captures rich mechanistic and mode-of-action information at the single-cell level while being at least 1000 times cheaper than other single-cell omics such as transcriptomics and proteomics, and at least 15 times cheaper than bulk -omics methods.

Prior work on inferring mode-of-action from Cell Painting and transcriptomics data in high-throughput, concentration-response toxicity screens has primarily relied on unsupervised approaches, where profiles from compounds with unknown modes-of-action are compared to groups with high-confidence annotations ^14–17^. Unsupervised methods are also commonly used to detect the concentration at which general bioactivity is observed in these screens ^18–20^. While these approaches have proven successful in many cases, they face limitations when compounds either perturb cells in multiple ways or trigger common compensatory responses, as subtler specific impacts can be masked by stronger generalized phenotypes. Supervised machine learning is an intuitive approach to disentangle specific mode-of-action and cell state signals from high-content omics data, and has successfully predicted toxicity-related outcomes from Cell Painting and transcriptomics data in single-concentration *in vitro* screens ^21–24^.

In this paper, we use supervised machine learning to predict diverse *in vitro* assay readouts relevant to mode-of-action and cell state using Cell Painting in primary human hepatocytes exposed to eight concentrations of 1085 compounds. These readouts include curated ToxCast/Tox21 assay activities as well as biochemical cytotoxicity assays measured alongside the Cell Painting data. We additionally investigate optimal strategies for making mode-of-action predictions from image-based profiles, including assessing different approaches for considering concentration and evaluating traditional and deep learning methods for extracting morphological features from images.

## Results & Discussion

### 1. High-throughput imaging detects compound bioactivity at lower concentrations than cytotoxicity assays

#### 1.1 Experimental design and data overview

Using the Cell Painting and two biochemical cytotoxicity assays, we tested 1085 compounds at eight concentrations ranging from 0.01 to 100 μM, with two replicates each, in primary human hepatocytes. The compounds were a subset of compounds from a list of 1495 compiled by the OASIS Consortium (Omics for Assessing Signatures for Integrated Safety) based on the public availability of *in vivo* hepatotoxicity data (**Supplementary Table 1**). The tested compounds include pharmaceuticals, agrochemicals, food additives, and known environmental contaminants. Hepatocytes were arrayed into 384-well plates and exposed to compounds for 44 hours before carrying out three assays. First, we took the supernatants from each well and measured lactate dehydrogenase (LDH), an enzyme that is released during cell membrane damage ^25^, and metabolic activity using the Realtime-Glo assay (Promega). Similar to the widely-used 3-(4,5-dimethylthiazol-2-yl)-2,5-diphenyltetrazolium bromide (MTT) assay which measures mitochondrial activity ^26^, Realtime-Glo measures substrate reduction by metabolically active cells, though it uses a luminescent rather than colorimetric readout. From hereon, we refer to Realtime-Glo assay as the MT assay to distinguish from the more common MTT assay.

Next, we applied Cell Painting dyes to stain the cellular DNA, RNA, mitochondria, actin, Golgi, and endoplasmic reticulum, and imaged each well at 40X magnification in five fluorescent channels. We created morphological profiles using three different computational methods, including CellProfiler software (yields single-cell level, named features for each combination of channel and cell compartment) ^27^, Cell Painting convolutional neural network (CP-CNN, pretrained on several large Cell Painting datasets, yields single-cell level, numbered features that are not directly interpretable nor mapped to channels) ^28^, and Meta’s “distillation with no labels” v2 vision transformer (DINO, pretrained on 142 million curated natural images, yields image-level, numbered features for each channel) ^29^. In addition to morphological features, we also counted the number of nuclei within each well using the Cell Painting images. We analyzed this cell count as a third cytotoxicity-related metric, alongside LDH and MT. While all three measurements – LDH, MT, and cell counts – reflect different biological aspects of cell state and toxicity, we collectively refer to them as “cytotoxicity-related” metrics for convenience.

#### 1.2 Concentration-dependent activity of compounds across different assay readouts

##### 1.2.1 40% of compounds induce activity as measured by LDH, MT, and cell counts

We analyzed the cytotoxicity-related endpoints to determine which compounds induced significant activity relative to DMSO negative controls. We also used concentration-response analysis to pinpoint the lowest concentration with detected activity for each active compound, herein referred to as the point-of-departure (POD), and whether the detected activity was more, or less, cytotoxic compared to DMSO. We detected activity for 430 of the 1085 compounds (40%) in the MT assay, 221 (20%) in the cell count assay, and 144 (13%) in the LDH assay for a total of 438 unique compounds. >98% of active compounds across all three assays increased cytotoxicity; very few decreased it. Overall, 429 compounds (40%) induced cytotoxicity as measured by one or more assays at one or more concentrations.

Most compounds active in the MT assay caused decreases in activity that are indicative of cytotoxicity, however ten compounds (1%) caused an increase in activity. Four of these compounds had particularly strong responses (**Figure 1A**), three of which (tolcapone, benzarone, and tiratricol) are pharmaceuticals with off-target impacts on the liver that have been mechanistically linked to the uncoupling of oxidative phosphorylation ^30–33^. Oxidative phosphorylation uncoupling decreases ATP production efficiency which causes a compensatory increase in the rate of oxidative phosphorylation, and so this could explain the observed increase in MT readouts for these compounds. The fourth compound is 2−ethylanthracene−9,10−dione (also called 2-ethylanthraquinone), an industrial compound used in the production of hydrogen peroxide. It exhibited both the highest potency and largest maximal response of all MT-increasing compounds in our study. According to the CompTox dashboard ^34^, between 50-500 tons of 2-ethylanthraquinone are produced or imported to the United States per year. It has over 100 related publications in material sciences journals but scant toxicological investigation besides its inclusion in several high-throughput screens ^10,35,36^. The ToxCast assay results summarized on the ‘Bioactivity’ tab of the CompTox dashboard provide evidence that 2-ethylanthraquinone could interfere with hepatocyte metabolism in similar ways as the other three compounds in **Figure 1A**. First, 2-ethylanthraquinone was a positive hit in twelve cell-based assays that assessed binding of eight different nuclear hormone receptors (RARA, NR1I2, ESR1, VDR, ESRRA, NR1I3, PGR, PPARG) at concentrations between 1-45μM and in the absence of cytotoxicity, which is consistent with our evidence that 2-ethylanthraquinone perturbs metabolism without causing cell death. Second, 2-ethylanthraquinone caused a decrease in mitochondrial depolarization at 14.8 μM, a cellular hallmark of oxidative phosphorylation uncoupling ^37^.

**Figure 1.**
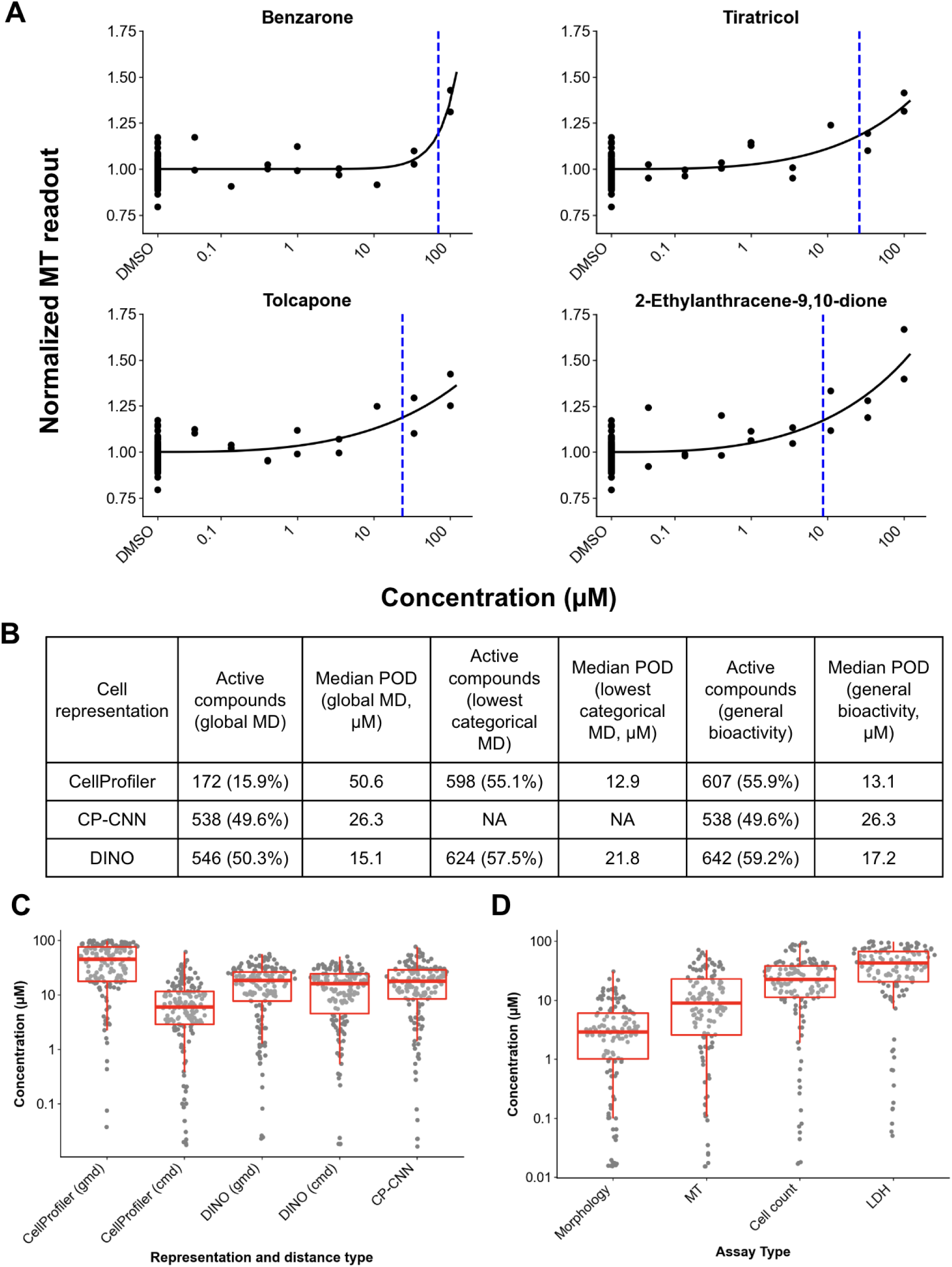
Compounds with a strong increase in MT readouts (A), with the POD indicated by blue, dashed lines. Number and percentage of active compounds and their morphological PODs according to different cell representations and distance metrics (B). Comparison of PODs across cell representations (C; n = 158, 15%) and assays (D; n = 131, 12%). In (C) and (D), only compounds with PODs for all four assays are displayed, to avoid bias for particular types of compounds. The morphological PODs in (D) are computed from CellProfiler categorical MDs. Lower values on the y axis indicate that activity was detected at a lower concentration (lower POD).

##### 1.2.2 Morphological profiling detects bioactivity for up to 60% of compounds

To analyze Cell Painting, we wanted to detect whether *any* impacts to morphology were observed for each compound, from hereon referred to as general bioactivity. To accomplish this, we summarized the image profiles using a Mahalanobis distance (MD)-based approach, as described previously ^18^. We computed the MD of each profile relative to the DMSO using all features (global MD) and using subsets of features (categorical MDs) corresponding to specific channels and/or cell compartments. Each of these distances is a measure of bioactivity, with larger values indicating that a profile has greater compound-induced perturbation of cell morphology. Next, we used concentration-response analysis as described for the cytotoxicity-related assays to determine PODs for each MD. We defined the general bioactivity PODs as the lowest POD among all of the statistically significant MD PODs for each compound.

Overall, morphological profiling and concentration-response analysis detected bioactivity for 16-59% of compounds, depending on the cell representation (CellProfiler, CP-CNN, and DINO) and distance metric used (global vs categorical) (**Figure 1B**). The two deep learning based methods (CP-CNN and DINO) detected ∼50-60% compounds as active, as did CellProfiler in the categorical MD case. The one outlier, at 16% active compounds, was CellProfiler using the global metric. Comparing the alternate approaches for extracting image features, CP-CNN general bioactivity PODs were, on average, 2.0-fold higher (less sensitive) than CellProfiler and 1.5-fold higher than DINO general bioactivity PODs (**Figure 1B and 1C**). There was no significant difference between CellProfiler and DINO general bioactivity PODs (paired t-test). However, DINO and CP-CNN general bioactivity PODs were more correlated with each other (r = 0.87, p-value < 1.0 e-14) than with CellProfiler (r = 0.70, 0.72; p-value < 1.0 e-14 for both DINO and CP-CNN).

#### 1.3 Morphological profiles are more sensitive than cytotoxicity readouts and align with assay biology

The morphological profiles from Cell Painting detected activity at lower concentrations than the three cytotoxicity-related readouts. There was a clear ranking of assay sensitivity, by both the number of active compounds and the relative POD, with morphology > MT > cell counts > LDH. The 131 compounds (12%) with PODs for all four assays qualitatively appear to be mainly conventionally toxic compounds. Among these, morphology PODs were 2.5-fold lower than MT, 7.9-fold lower than cell counts, and 15.8-fold lower than LDH (Figure 1C, all p-values < 1.0 e-14). The assays’ differences in sensitivity are consistent with assay biology. LDH, an enzyme released during membrane damage, is more representative of the subset of cell death pathways that cause catastrophic membrane rupture, for example necrosis, and represents a relatively late stage of cell death. Cell count represents the balance between all forms of cell proliferation and cell death, so we expect it to be active for more compounds and at lower concentrations than LDH. The MT assay measures metabolic competence, which we expect to be perturbed at lower concentrations than those that cause outright cell death. Finally, Cell Painting captures subtle morphological changes across a wide biological space far beyond cell death alone, having been shown to detect a wide range of chemical and genetic perturbation modes-of-action ^24,38^.

### 2. Morphological profiles predict exposure-level bioactivity in paired cytotoxicity assays

#### 2.1 Morphological profiles predict cytotoxicity assay readouts

The MT and LDH assays measure different aspects of cytotoxicity; we wanted to know whether Cell Painting better captured these nuances than simple cell counts and technical factors such as well position that are known to influence cell viability. To assess this, we trained XGBRegressor models to predict MT and LDH readouts based on Cell Painting profiles (CellProfiler, CP-CNN, and DINO representations), and compared their performance to several baseline models.

First, there were no significant differences in performance across representations (**Table 1**), thus we refer to morphology-based predictions from all representations as the “Cell Painting” predictions from hereon. We found that Cell Painting outperformed the cell count and technical baseline for the MT assay but not for the LDH assay (**Table 1**). To some degree, the cell count captures the same types of cytotoxicity captured by the LDH assay. As well, there is also a strong influence of technical plate and well position factors on the LDH readouts (**Supplementary Figure 1**), which resulted in high variability between technical replicates. In fact, the R^2^ between observed LDH readouts and predictions based on either Cell Painting or the technical baseline models were significantly higher than the R^2^ between technical replicates of the same compound and concentration (mean difference = 0.20, p-value < 1.0 e-14). The MT readouts were less influenced by well position, showed much better concordance across replicates, and were better predicted by Cell Painting than by the technical baseline.

**Table 1.** Mean R^2^, root mean square error (RMSE), and mean average error (MAE) between predicted and observed biochemical assay readouts for models trained on different input features across 10 different train-test splits (by compound). Standard deviation values are in parentheses. The technical baseline includes features for cell count, batch, plate, and well position. The replicate baseline computes metrics between the first and second replicate of each compound-concentration exposure and captures the consistency between replicates for that assay. The mean predictor baseline predicts the mean assay value for all compounds, providing a minimum performance reference that any useful model should exceed.

| Input features | $R^2$ | RMSE | MAE |
| --- | --- | --- | --- |
| LDH (ridge-normalized) |  |  |  |
| Mean predictor baseline | -0.002 (0.0020) | 0.1 (0.011) | 0.061 (0.0030) |
| Technical baseline | 0.65 (0.044) | 0.06 (0.0049) | **0.035 (0.00078)** |
| Replicate baseline | 0.45 (0.062) | 0.08 (0.0025) | 0.053 (0.0017) |
| CellProfiler | 0.65 (0.051) | 0.059 (0.0026) | 0.039 (0.00085) |
| CP-CNN | **0.66 (0.053)** | **0.058 (0.0025)** | 0.039 (0.0010) |
| DINO | **0.66 (0.052)** | 0.059 (0.0030) | 0.04 (0.0011) |
| MT (ridge-normalized) |  |  |  |
| Mean predictor baseline | -0.0019 (0.0017) | 0.17 (0.014) | 0.092 (0.0044) |
| Technical baseline | 0.70 (0.055) | 0.093 (0.0050) | 0.048 (0.0014) |
| Replicate baseline | **0.88 (0.020)** | **0.06 (0.0052)** | **0.032 (0.0013)** |
| CellProfiler | 0.79 (0.039) | 0.077 (0.0034) | 0.042 (0.0013) |
| CP-CNN | 0.79 (0.040) | 0.077 (0.0032) | 0.042 (0.0011) |
| DINO | 0.79 (0.039) | 0.078 (0.0031) | 0.042 (0.0013) |

#### 2.2 Prediction outliers reveal biologically meaningful differences between morphology, cell counts, and cytotoxicity-related assays

We wanted to know which additional biological signals were captured by Cell Painting compared to the technical and cell count baseline for the MT predictions. We used publicly available protein target annotations of compounds and overrepresentation analysis to determine whether any targets were enriched among the compounds for which the Cell Painting MT predictions were significantly better (*n* = 138, 1.0%) or significantly worse (*n* = 304, 2.3%) than the technical baseline predictions. Exposures whose MT values were better predicted by Cell Painting were significantly enriched in 480 molecular targets (**Supplementary Table 2**), many of which are in the cell cycle (29/97 targets, FDR = 1.46e-12), PI3K-Akt signaling (53/256 targets, FDR < 1.0 e-14), MAPK signaling (47/249, FDR = 1.90e-12), and p53 signaling (21/64 targets, FDR = 3.43e-10) KEGG pathways. Exposures that were better predicted by Cell Painting tended to have lower normalized MT values compared to those better predicted by the technical baseline and compared to all samples (Figure 2A). The cell counts were also lower, although most were in a normal range between 500 and 1000 (Figure 2B). We wanted to know whether the samples better predicted by Cell Painting had one or multiple distinct phenotypes, however clustering of compounds based on pairwise cosine similarity between samples was mainly dominated by differences in cell count (Figure 2C), highlighting the difficulties of using unsupervised approaches to analyze toxic exposures. Because Cell Painting encodes cell counts and technical effects in addition to interesting biological signals ^39^, we expected that the samples with MT readouts better predicted by the technical baseline compared to Cell Painting would be random and have no meaningful biological signals. There were in fact no significantly enriched targets in this list of samples (**Supplementary Table 3**), increasing confidence that target enrichment analysis is a valid approach.

**Figure 2.**
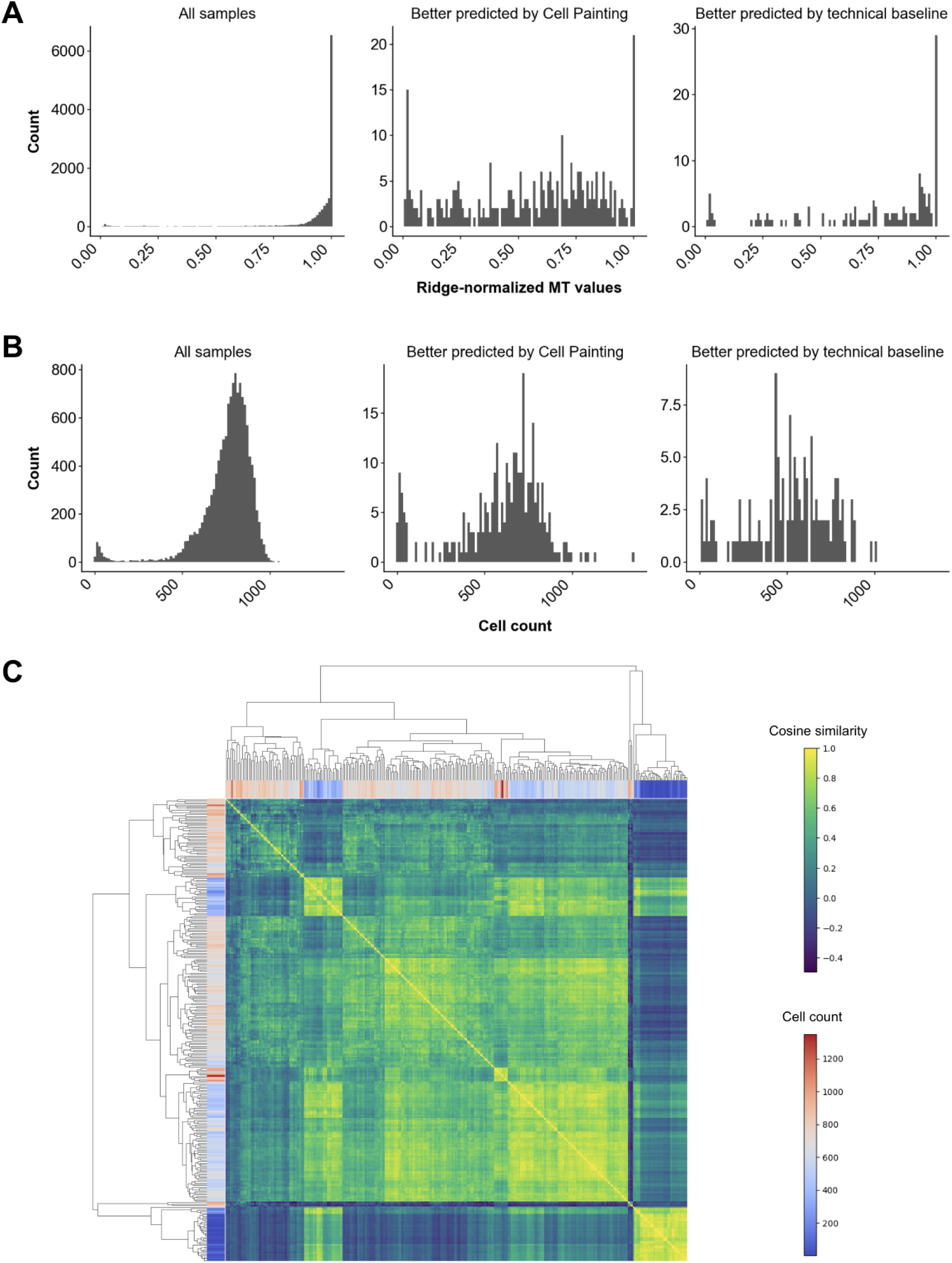
Ridge-normalized MT values (A) and cell counts (B) across different sets of samples. (C) Clustergram of pairwise cosine similarity between DINO profiles of the exposures that were significantly better predicted by Cell Painting than by the technical baseline, with rows and columns colored by cell count.

Like any regression problem, our analysis revealed cases where predicted values deviated from observed measurements. We leveraged these prediction discrepancies to investigate the biological mechanisms underlying the relationship between cellular morphology and metabolic activity. Specifically, we found that a small number of wells had MT readouts that were poorly predicted by both their morphological profiles and technical features like cell count, suggesting fundamental disconnects between cellular appearance and metabolic activity.

We analyzed the exposures with significantly higher morphology-predicted MT readouts than observed (*n* = 261, 2.0%), with target enrichment analysis to investigate whether there were any particular modes-of-action underlying this phenomenon, and found that they were enriched in compounds targeting 147 proteins (FDR < 0.05, **Supplementary Table 4**). These targets included many cytochrome P450 enzymes, for example CYP3A5, CYP3A43, CYP2B6, CYP3A7, CYP2C8, and CYP3A4. They also included many compounds that targeted G-protein coupled dopaminergic, serotonergic, and adrenergic receptors (amoxapine, apomorphine, aripiprazole, cisapride, clomipramine, domperidone, paliperidone, risperidone, and tamsulosin) - this is interesting because neurotransmitters are known to regulate hepatic CYP expression and metabolism ^40–42^. The MT assay specifically measures mitochondrial dehydrogenase activity, which depends on the flow of electrons through NADH and FADH2 to the electron transport chain ^26^. We hypothesize that CYP induction and general metabolic reprogramming could decrease MT values without a dramatic change in morphology or cell count. First, compound exposures could shift metabolism from oxidative phosphorylation to other pathways such as glycolysis that do not require mitochondrial activity ^43^. In this case, hepatocytes would be metabolically active with morphological profiles indicative of metabolic competence, and yet have low rates of MT substrate reduction. Mitochondrial activity could be further reduced if the metabolic perturbations increase the reactive oxygen species (ROS) burden within the cells, which can directly impair the dehydrogenase enzyme activity required for MT substrate reduction ^44,45^. The overpredictions were also enriched for compounds targeting xenobiotic transporters including ABCB1, ABCG2, SLCO1B1, and SLCO1B3. If the MT substrate partially relies on xenobiotic transporters to enter cells, then compounds that inhibit these transporters could lead to lower MT readouts simply due to lower levels of MT substrate within cells.

Similarly, analyzing the underpredicted cases (n = 131, 1.0%) revealed enrichment for compounds targeting distinct biological processes, including proteasome inhibition (specifically through bortezomib, carfilzomib, and ixazomib exposures), xenobiotic metabolism, bile acid synthesis, general cell stress, and apoptosis (**Supplementary Table 5**). Some of these underpredictions might be explained by assay interference by the administered compounds. For example, redox-active compounds can directly reduce the MT substrate and some compounds have a similar absorbance spectrum to the reduced substrate ^46,47^, leading to higher MT assay readouts than supported by the actual level of mitochondrial activity.

### 3. Morphological profiles predict activity in external assays, depending on relevance of experimental factors

#### 3.1 Preparing ToxCast assay endpoints

We wanted to assess whether Cell Painting morphology profiles from human hepatocytes could predict a broader set of toxicity-related assay outcomes beyond MT and LDH. ToxCast is a rich source of targeted assay data for a set of compounds that includes 963 (89%) of our tested compounds. We curated 48 cytotoxicity endpoints, 292 non-cytotoxicity cell-based endpoints, and 72 cell-free endpoints. Cytotoxicity endpoints were measured alongside many of the more specific ToxCast endpoints, resulting in many repeated cytotoxicity readouts for the same compound and often the same cell type. As a result, benchmarking assay prediction would over-represent simple cytotoxicity endpoints at the expense of more useful and specific toxicity endpoints. We therefore curated cytotoxicity endpoints to be consensus hit calls of cytotoxicity readouts aggregated across 28 cell types and 20 tissue sources resulting in 48 (consensus-level) readouts. While the cytotoxicity hit calls broadly agree across different cell and tissue types, there are distinct clusters of compounds with cytotoxicity in some cell types but not others (**Supplementary Figure 2**).

Most of the cell-based endpoints were measured in cell types derived from liver (52%), vascular tissues (22%), and kidney (12%). They fall into six different categories, with 84% assessing the activity of individual molecules, such as mRNA transcript levels or activation/inhibition of specific receptors, and the remainder assessing perturbations at the pathway, subcellular, or cellular levels. The individual mRNA and protein targets come from 36 distinct protein families, with the three most common being nuclear receptors (19%), DNA binding proteins (18%), and cytokines (11%). The cell-free endpoints assess two main functional categories: protein binding (51%) and enzymatic activity (49%). The cell-free targets come from 11 protein families, with the three most common being GPCRs (29%), CYP450s (15%), and nuclear receptors (14%). Overall, the median number of tested compounds was 346 for cytotoxicity endpoints, 306 for cell-based endpoints, and 33 for cell-free endpoints, and the median percentage of active compounds was 21% for cytotoxicity, 7% for cell-based, and 41% for cell-free (**Supplementary Figure 3**). Many of the positive hits in the cell-free assays were for the enzymatic activity of various cytochrome P450s, which are expected to interact with many xenobiotic compounds.

#### 3.2 Cell Painting profiles predict activity in ToxCast assays

Cell Painting profiles predict compound activity in cytotoxicity and cell-based assays, but not cell-free assays, compared to a random baseline (Figure 3). We determined this by converting ToxCast data into binary hit calls of “active” or “inactive” for each compound-endpoint pair, median-aggregating CellProfiler features across all concentrations’ consensus profiles for each compound, and training a suite of binary XGBoost classifiers to predict compound activity for each endpoint based on the consensus profiles. Model performance was assessed with AUROC, which measures the ability of the model to distinguish between active and inactive compounds across all possible classification thresholds, and PRAUC, which measures the precision-recall tradeoff and emphasizes performance on the active class where data may be imbalanced ^48^. We also converted the paired LDH and MT readouts into binary hit calls so that we could benchmark the ToxCast prediction performance against assays with an experimental design that is perfectly aligned with the Cell Painting profiles. Cell Painting profiles performed best at predicting cytotoxicity assays, including the MT and LDH assays; this is not surprising given that the image profiles include cell count in addition to morphological changes associated with cell stress and death. Cell-based assays were next-most predictable, with cell-free assays being least, in line with Cell Painting being itself a cell-based assay. Assay activity prediction based on cell count alone outperforms the random baseline for cytotoxicity but not cell-based or cell-free assays (Figure 3).

**Figure 3.**
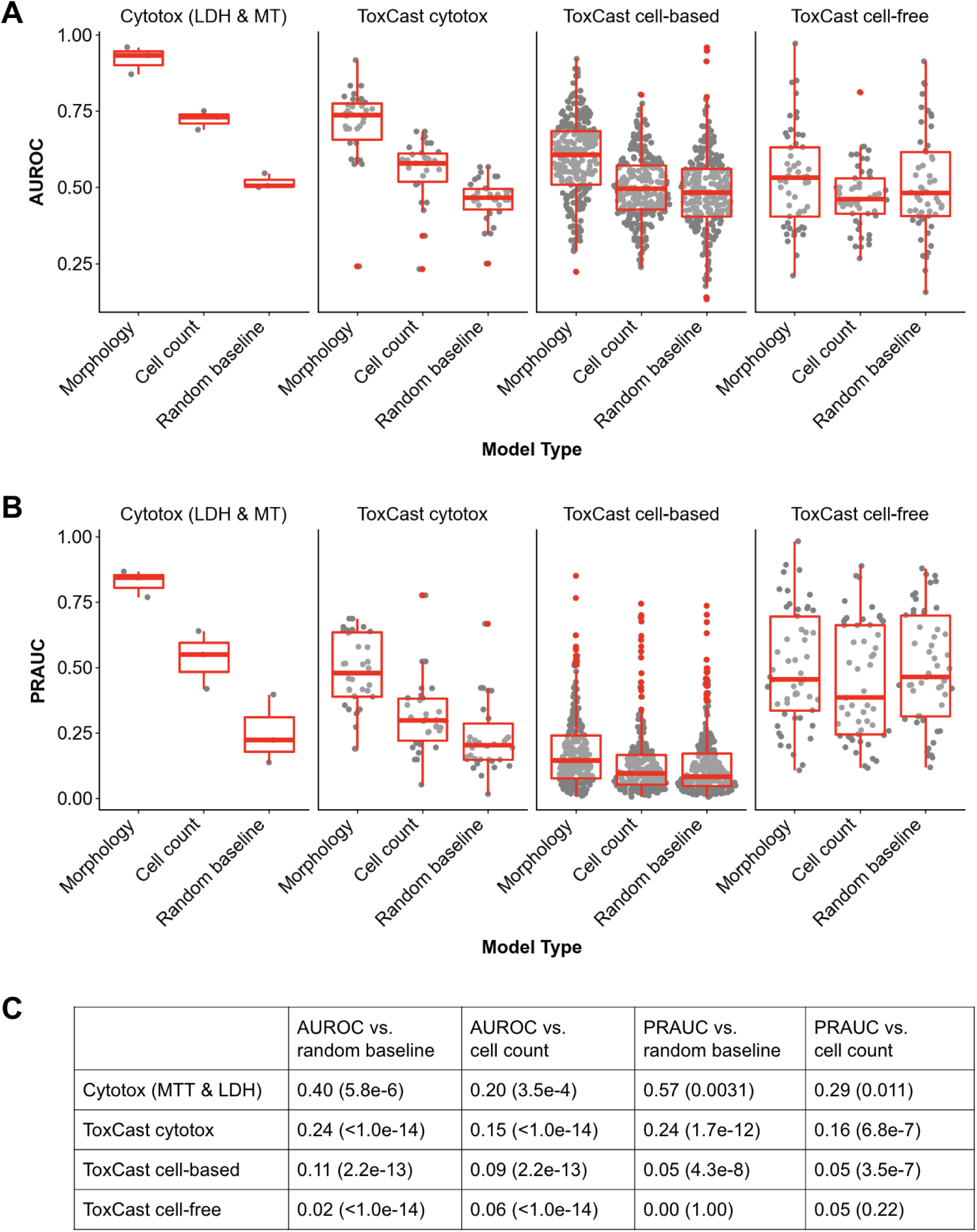
Distributions of AUROC scores (A) and PRAUC scores (B). The morphology and cell count classifiers are trained on CellProfiler features and cell counts that were median-aggregated across all concentrations. The random baseline is trained on CellProfiler morphology features after randomly shuffling the labels. (C) Mean difference in classifier performance according to AUROC and PRAUC, stratified by endpoint type, compared to random and cell count baselines. Associated paired t-test p-values are in parentheses.

##### 3.2.1 Consensus profiles that contain all concentrations are simple and performant

We hypothesized that filtering out Cell Painting profiles from non-bioactive concentrations would be more predictive of assay activity than including them. To test this, we created a second set of consensus profiles for each compound using only profiles from concentrations greater than the general bioactivity POD, and used these to predict assay activity (Figure 4A). Interestingly, there was only a small improvement in the AUROC scores for the bioactive-concentration versus all-concentration consensus profiles for the ToxCast cytotoxicity (mean difference = 0.04, FDR < 1.0 e-14) and ToxCast cell-based (mean difference = 0.02, FDR = 0.002) assays. There were no significant differences in AUROC for the other assays, nor for the PRAUC scores for any assays (**Supplementary Figure 4A**). We next hypothesized that additionally filtering out exposures that induced cytotoxicity would improve predictive performance for the targeted cell-based and cell-free assays because this would remove non-specific signals associated with general cell stress and death pathways, though this might be offset by the reduction in useful data points. To test this, we created a third set of consensus profiles using only concentrations greater than the morphological POD and lower than the cell count POD. As expected, filtering out the cytotoxic profiles resulted in significantly worse performance for both of the cytotoxicity assay categories (FDR < 1.0 e-14 for both AUROC and PRAUC). Contrary to our prediction, there were no significant differences in performance for the bioactive-but-not-cytotoxic consensus profiles compared to the bioactive profiles for the cell-based and cell-free assays.

**Figure 4.**
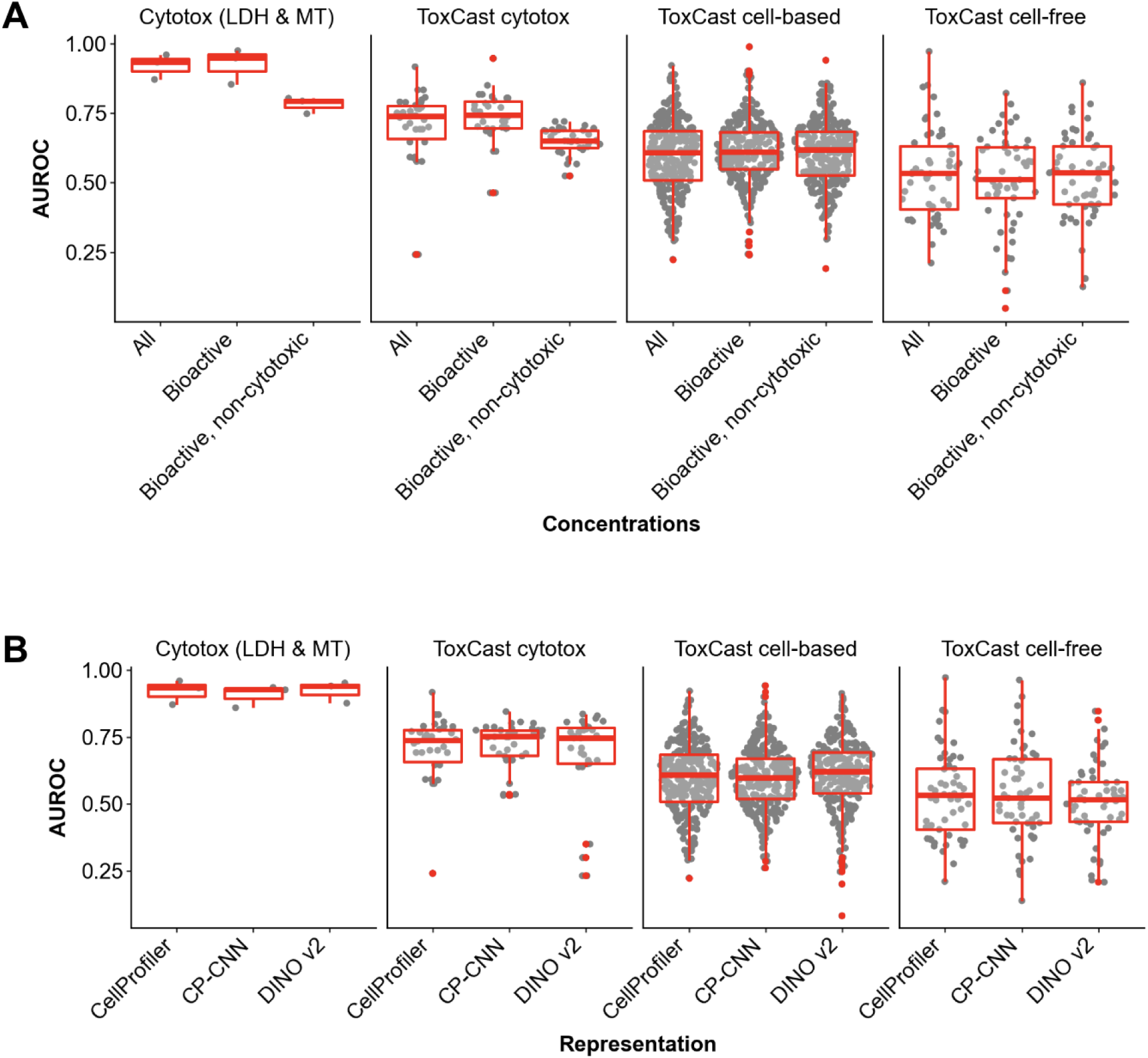
Classifier AUROCs, stratified by endpoint type, across consensus profile strategies (CellProfiler features) (A) and cell representations (“all” concentration consensus profiles) (B).

One possible explanation for the all-concentration profiles generally performing at or near optimal is that plate and well-position effects are more completely “averaged out” across the higher number of profiles used to create the all-concentration consensus profiles, and that this benefit outweighs the benefits from boosting the compound-induced signals. Another explanation is that defining morphology-based bioactivity PODs based on Mahalanobis distances is not very sensitive and consistently overestimates bioactivity PODs, such that relevant information is lost when they are used as filtering thresholds. This is consistent with previous findings that Cell Painting Mahalanobis distance-based bioactivity was detected at consistently higher concentrations than those that caused activity in targeted ToxCast assays ^49^. We therefore predict that the benchmark presented in Figure 4A could be used to evaluate new approaches for computing bioactivity PODs from Cell Painting data - an open challenge for the more general toxicogenomics community ^20^. Computational methods with higher sensitivity and specificity for bioactivity detection from high-dimensional omics profiles should outperform the all-concentration and bioactive-concentration consensus profiles used here. Given the current small, if any, performance increase associated with defining custom concentration ranges, we recommend that aggregating all concentrations into a consensus profile is a simple yet performant approach for this type of supervised analysis.

##### 3.2.2 Alternative cell representations perform similarly

Although there is great enthusiasm for using deep learning strategies to extract features from raw pixels in images, the predictive performance was very similar across the different cell representations we tested. DINO had slightly higher AUROC scores for the cell-based ToxCast assays (mean diff = 0.02 for both comparisons, p-value = 2.7e-5 and 7.4e-6 for CellProfiler and CP-CNN respectively); otherwise there were no significant differences in performance according to AUROC or PRAUC scores (Figure 4B, **Supplementary Figure 4B**).

#### 4. Discussion

Overall, we found that morphological profiles from Cell Painting in human hepatocytes detect bioactivity at lower concentrations than multiple assays of cytotoxicity. They also have some ability to predict many mode-of-action and toxicity-related assay endpoints in cell-based assays, but not cell-free assays of protein binding and enzymatic activity. These findings demonstrate the potential of morphological profiling as a sensitive, early-stage screening tool for compound toxicity.

In this analysis, state-of-the-art deep learning features had very similar performance to traditional engineered image features from CellProfiler. This has not been an unusual finding in the literature but could reflect non-optimal feature extraction; there are many alternatives one could try ^50,51^. Beyond predictive performance, the different cell representations we tested had relative strengths and limitations in their practical implementation. DINO was the easiest end-to-end solution because it was used off-the-shelf and did not require any cell or nucleus segmentation; however this also means that it did not produce profiles at the single-cell level, which might be desirable for some applications ^52,53^. CP-CNN was also relatively hands-free, with only a single tunable parameter (CellPose nuclei size). CellProfiler, by comparison, requires tuning in two hands-on steps (nuclear and cell segmentation) each with dozens of potentially tunable parameters, though pipelines, once expertly tuned, can often be reused on similar data with little to no further tuning. However, the standard CellProfiler processing pipeline yields readily interpretable features and includes standardized QA/QC metrics, which have proved invaluable in our past experience. We highly recommend that similar quality checks be developed for deep learning cell representation pipelines.

The ability to predict multiple targeted assay endpoints from a single morphological profile supports the cost-effectiveness of profiling approaches compared to running hundreds of individual assays ^4^. However, image-based profiles are less immediately biologically interpretable relative to targeted assays for toxicity, particularly those measuring key events in adverse outcome pathways (e.g., apoptosis, ER stress, or estrogen production). The interpretability challenge is especially pronounced for high-content imaging, which is inherently less interpretable than omics approaches that measure discrete molecular features, such as transcriptomics or proteomics. Our findings demonstrate that Cell Painting profiles do contain rich mode-of-action signals and can discriminate subtle differences in cytotoxicity, given sufficient high-quality training data. However, the ability to detect robust signals was related to how closely the targeted assay experimental design aligned with Cell Painting assay design. ToxCast endpoints were measured in a wide range of assay types, including cell-free, cell-based from many different cell types, tissue-based, and *in vivo*. Compound exposures were conducted at different concentration ranges and time points, and some assays included additional environmental factors, for example co-exposure with lipopolysaccharide to assess immune function.

Future work could advance the utility of imaging and omics data in compound toxicity screening. Publicly available training data (both from targeted assays and *in vivo* exposures) is precious and every effort should be made to document the experimental design parameters discussed above, harmonize units and other nomenclature, and improve the translation between data sources via pharmacokinetic and chemical partitioning models ^54,55^. Given the relative scarcity of well-annotated, targeted training data, comparisons are often made across datasets collected for different cell types, and it is unclear how much this impacts results. While this study tested the ability to predict outcomes that were measured in different cell types, it was challenging to derive clear conclusions because the assay endpoints measured in each cell type were also different. Future work by the OASIS Consortium will collect Cell Painting data for additional cell types for a consistent set of compounds, enabling direct comparisons of how well cell types with diverse lineages predict the same targeted assays. For a subset of compounds and cell types, it will also capture transcriptomics and proteomic data, enabling a comparison and combination of these data types with imaging, which may prove powerfully predictive.

## Acknowledgements

This work was supported by Banting and SOT Syngenta postdoctoral fellowships to JE, a National Institutes of Health grant (NIGMS R35 GM122547 to AEC), and the Omics for Assessing Signatures for Integrated Safety (OASIS) Consortium, which is supported by a grant from the Massachusetts Life Sciences Center (MLSC) Bits to Bytes Capital Call grant to SS, as well as partnered resources and expertise from industry partners and in-kind contributions from academic, government, non-governmental organizations, and biotech partners (listed here https://oasisconsortium.org/members)

## Code & Data Availability

Images and profiles are available here: https://cellpainting-gallery.s3.amazonaws.com/index.html#cpg0037-oasis/axiom/

The codebase is available here: https://github.com/broadinstitute/2024_09_09_Axiom_OASIS

## Competing Interests

The Authors declare the following competing interests: B.A.C., S.S., and A.E.C. serve as scientific advisors for companies that use image-based profiling and Cell Painting (B.A.C.: Marble Therapeutics, A.E.C: Recursion, SyzOnc, Quiver Bioscience, S.S.: Waypoint Bio, Dewpoint Therapeutics, Deepcell) and receive honoraria for occasional scientific visits to pharmaceutical and biotechnology companies. K.J.K. is a consultant for Tome Biosciences, AlloDx, and Vor Biosciences, and a member of the scientific advisory board of Nurture Genomics. All other authors declare no competing interests.

## Methods

### Curation of Compounds

The OASIS Consortium curated a total of 1495 compounds known to be of interest to the hepatotoxicology community by integrating data from established toxicological resources, including DrugMatrix, TG-GATES, ToxRefDB, DILIlist and DILIrank ^56–60^. The curated dataset is released via https://oasisconsortium.org/oasis-compounds. Compounds were cross-referenced using InChIKey identifiers to remove duplicates. The finalized list was a subset of 1085 compounds that could be sourced from MedChemExpress.

### Cell culture

384-well microplates (Corning #4518) were coated with 50 µg/mL of Collagen I, Rat Tail (Thermo Fisher) diluted in 20 mM acetic acid. Plates were incubated with the collagen solution for 1 hour, washed twice with sterile H₂O, and then air-dried in a sterile cabinet for a minimum of 2 hours before cell seeding. Pooled primary human hepatocytes (PHH) from BioIVT (Liverpool 5-donor pool) were thawed in a 37°C water bath following the vendor’s protocol, resuspended in INVITROGRO CP medium supplemented with TORPEDO antibiotic mix (Complete CP medium), and plated at 5,500 cells/well. Plates were incubated at 37°C in a humidified 5% CO₂ incubator (Cytomat) for 3–5 hours to allow time for cells to become adherent to the collagen coating.

### Compound treatment

Once cells were attached, the medium was replaced with Complete CP medium containing test compounds at a 2x final concentration. Plates were then returned to the incubator for 44 hours at 37°C, 5% CO₂, until the assay endpoints were measured.

### LDH and MT biochemical assays for cytotoxicity assessment

LDH (CyQUANT™ LDH Cytotoxicity Assay, Invitrogen) and MT (RealTime-Glo MT Cell Viability Assay, Promega) assays were performed to assess cytotoxicity. For the LDH assay, a clear microplate (Greiner 781101) was pre-filled with 7.5 µL HBSS, and 2.5 µL of cell media was transferred from the cell plate to the clear LDH assay plate. The assay was performed according to the manufacturer’s instructions by adding 10 µL of reaction mixture, incubating for 30 minutes in ambient conditions, then adding 10 µL of stop solution before measuring absorbance at 530 nm and 720 nm on an EnVision plate reader. For the MT assay, the cell viability substrate and NanoLuc® enzyme were added to pre-warmed Complete CP medium to prepare the reagent solution, which was added at 15 µL/well to the cell plate and incubated for 1 hour at 37°C and 5% CO₂ before reading luminescence on the EnVision plate reader. Cell plates were washed 2x with HBSS to remove any residual luminescent substrate prior to staining with Cell Painting dyes, which could interfere with image-based readouts.

### Cell Painting and image acquisition

We used the Cell Painting v3 protocol (Cimini et al., 2023) using PhenoVue Cell Painting JUMP Kit (Revvity, PING21) to generate fluorescent images. Initially, PhenoVue 641 Mitochondrial Stain in CP complete medium was added at a final concentration of 500 nM to HBSS-washed cells as described above, and cells were incubated for 30 minutes at 37°C, 5% CO₂. Cells were then fixed with 4% paraformaldehyde (PFA) for 20 minutes at room temperature and washed four times with HBSS. The remaining dyes, including PhenoVue Fluor 488 Concanavalin A (5 µg/mL), PhenoVue 512 Nucleic Acid Stain (1 µg/mL), PhenoVue Fluor 555 WGA (1.5 µg/mL), and PhenoVue Fluor 568 Phalloidin (8 nM), were added. We reduced the concentration of PhenoVue 512 Nucleic Acid Stain (RNA) from 3 µg/mL to 1 µg/mL to reduce channel similarity to the neighboring WGP channel; otherwise, the protocol was followed as published. Images were acquired on an Operetta imaging system (Revvity) using a 40x water objective in non-confocal mode, sampling 15 fields of view (FOVs) per well. Full-resolution images (2160×2160) were captured with binning set to 1. A single z-plane was acquired for each of the five fluorescent channels and one brightfield channel, with the optimal z-plane selected for each channel based on focal quality assessed by maximal variance in laplacian texture measurement.

### Image processing

#### CellProfiler

We used CellProfiler bioimage analysis software (version 4.2.4) ^27^ to process the images using classical algorithms following a prior protocol ^61^. Flat field correction was applied to the images ^62^. We segmented nuclei using Cellpose (version 2.3.2) ^63^ as a CellProfiler plugin^64^ with the pre-trained “nuclei” model and minimum object size of 1500 px) and cells and measured feature categories including fluorescence intensity, texture, granularity, density, and location (see http://cellprofiler-manual.s3.amazonaws.com/CellProfiler-4.2.4/index.html for more details) across all segmented compartments and all imaged channels. We obtained 5640 feature measurements from each of about 800 cells per well. We parallelized our image processing workflow using Distributed-CellProfiler ^65^. The image analysis pipelines we used are available in the Cell Painting Gallery ^66^.

#### CellPainting-CNN (CP-CNN)

Images were converted to JPEG-XL format with the -q 99 option. Cell centers were taken from CellProfiler output. DeepProfiler (version bed9d6a with pull request 358) with the CellPainting-CNN model ^28^ [https://github.com/cytomining/DeepProfiler/pull/358/files, https://zenodo.org/records/7114558] was used to generate embeddings.

#### DINO

Each channel image was preprocessed with instance normalization at the field level to control for intensity variation. Then, a pre-trained DINOv2 model ViT-L/14 (https://github.com/facebookresearch/dinov2) was used on each of the 6 normalized channel images per FOV to generate vectors of features of length 768 per channel.

### ToxCast curation

ToxCast includes data from different assays, each of which can have one or multiple endpoints, measured across different tissue, cell, and cell-free models from different species. Each endpoint was measured at multiple compound concentrations, often along with cell viability or cytotoxicity endpoints run in parallel to the assay. For each compound-endpoint concentration-response series, the ToxCast authors produce a hit call from 0 to 1 (with 1 being high confidence that there is a change of activity relative to the control), and an AC50 value (concentration at which 50% of the maximum endpoint value is observed) ^11^. We matched ToxCast compounds to our tested compound library using the EPA’s DTXSID.

We accessed all assay endpoint data from *invitrodb* version 4.1 using the SQLalchemy version 1.4.54 Python package. We filtered endpoints to only include those from human cell and cell-free assays. We excluded QA/QC endpoints like background fluorescence, the individual ch1 and ch2 endpoints used to compute the more interpretable “ratio” endpoints, any assay marked as “follow-up”, and any human cells that were transfected with non-human genes. When the same endpoint was measured after multiple exposure time points, we kept the single timepoint that most closely matched our experimental design (44 hour compound exposure). Any positive hit call with an associated AC50 higher than 100uM (the highest concentration we tested) was set to 0.

A significant proportion of the specific endpoint AC50s are at the same or higher concentrations than where cytotoxicity is observed; these are more indicative of general cell stress than of the specific modes-of-action that the endpoints were intended to measure ^67,68^. We computed “consensus cytotoxicity hit calls” across different biological categories defined by cell line and tissue annotations. For each category, we considered a compound to have a positive cytotoxicity hit call if 20% or more of the individual hit calls were positive and defined the AC50 as the median AC50 of all of the positive hit calls. Next, we compared each specific endpoint AC50 to the closest matched consensus cytotoxicity AC50 (cell-level if available, tissue-level otherwise). If the specific AC50 was higher than the cytotoxicity AC50 / 2, then the specific endpoint hit call was set to 0. Across all cell-based assays and OASIS compounds, there were 281,196 hit calls for non-cytotoxicity endpoints, where each hit call specifies whether a particular assay was active for a particular compound. Of these, there were initially 17,110 positive hits (6.1% of all hit calls) and filtering for cytotoxicity-confounded hits reduced the positive calls by 51% from 17,110 to 8400 (3.0% of all hit calls). Finally, we removed endpoints from the curated dataset if there were fewer than five positive and negative hit calls in our tested compound library.

### Profile processing

Profiles from CellProfiler, DINO, and CP-CNN were processed with the same pipeline. First, features with missing or infinite values were filtered out. Next, features that had an absolute coefficient of variation < 0.001 in the DMSO negative control samples on any plate were also filtered out to prevent the creation of exploding values in the next step. Profiles were next MAD-normalized, by subtracting the median and dividing by the MAD of the DMSO samples from the same plate, to minimize systematic plate effects. Finally, profiles were filtered to remove low variability and highly correlated features using the “variance_threshold” and “correlation_threshold” from Pycytominer (v1.2.0) using default parameters.

### Mahalanobis distance calculations

Various Mahalanobis distances were computed between each morphological profile and the centroid of the DMSO samples on the same plate, as described previously ^18^. The global Mahalanobis distance is calculated using entire profiles that include all measured features. The categorical Mahalanobis distances are computed for “mini” profiles, each containing a subset of related features. The set of categories depends on the type of cell representation.

CellProfiler produces named, interpretable features and so it is possible to group them into categories according to both the subcellular compartments imaged by each channel (DNA, ER, RNA, actin+golgi, and mitochondria) and the various segmented parts of each cell (entire cell, cytoplasm only, and nucleus only) ^27^. In addition to the five channels, we also considered the “AreaShape” features as a sixth category since these do not depend on any individual channel. This resulted in 6*3 = 18 categories, for example Cells_RNA or Cytoplasm_AreaShape.

Cell representations from deep learning methods do not have features with meaningful names. Since the CellPainting-CNN produces one set of embeddings that represent images from all channels simultaneously ^28^, it is not possible to break these into smaller categories and so only global Mahalanobis distances were computed for this representation. The DINO architecture produces one set of embeddings for each individual image, and so we computed six categorical Mahalanobis distances for each perturbation, one for each channel (DNA, ER, RNA, actin+golgi, mitochondria, and brightfield).

### Concentration-response analysis

Curve fitting was performed on all Mahalanobis distances (both global and categorical, where applicable), cell count, and normalized MT and LDH biochemical assays using fastbmdR (v0.0.0.9, https://github.com/jessica-ewald/fastbmdR)^69^. Prior to fitting, concentrations were converted to a logarithmic scale (base 10). Eight parametric models (Exp2, Exp3, Exp4, Exp5, Poly2, Lin, Power, Hill, as defined here ^19^) were fit to the Mahalanobis distances, cell count, and biochemical assay readouts for each compound. Curve fits were filtered out if the standard deviation of the residuals was more than three times the standard deviation of the DMSO controls. The best-fit model for each readout for each compound was defined as the one with the lowest standard deviation of the residuals. The POD of each best-fit model was determined as the concentration at which the fitted curve surpassed the 95th percentile of the DMSO samples. PODs were filtered out if the ratio between the upper and lower 95th confidence intervals was greater than 40 or if the BMD was higher than the highest tested concentration. For morphology readouts, compound-level PODs were defined as the lowest POD among all of the global and categorical Mahalanobis distance PODs that passed the QA/QC filters for each compound.

### Supervised analysis

XGBoost was used to predict various continuous and categorical labels from different sources^70^. For all scenarios, five-fold stratified cross-validation with compound splits and default parameters were used to assess general predictive performance. For continuous outcomes, we trained XGBRegressor models and compared performance to a mean predictor and to a suite of baselines that used cell count and various technical metadata such as source, plate, and well position. We assessed performance with R^2^ and RMSE and MAE. For the categorical outcomes describing assay activity, we trained binary XGBoost classifiers and compared performance to a random baseline (labels shuffled prior to training) and to a cell count baseline. Classification performance was assessed using AUROC and PRAUC.

The continuous outcomes had one readout per Cell Painting profile, therefore we trained models based on individual profiles. The categorical outcomes describing assay activity had one label for each compound, corresponding to sixteen morphological profiles (two replicates for each of the eight tested concentrations). We created three different compound consensus profiles computed as the median of each feature across different sets of profiles: 1) all profiles (“all”), 2) all profiles after the morphology POD, defaulting to all profiles if there was no detected morphological change (“all_morph”), and 3) all profiles between the morphology POD and cytotoxicity POD, defaulting to all profiles after the morphology POD if there was no cytotoxicity and to all profiles if there was no cytotoxicity or morphology changes (“all_morph_cytotox”). We trained a different classifier for each consensus profile, and compared performance across consensus profile strategies.

## Notes

https://github.com/broadinstitute/2024_09_09_Axiom_OASIS

https://cellpainting-gallery.s3.amazonaws.com/index.html#cpg0037-oasis/axiom/

