## Supplementary Figures for "Cell Painting for cytotoxicity and mode-of-action analysis in primary human hepatocytes"

**S1** - Plate effects of cell count, LDH, and MT
**S2** - ToxCast cytotoxicity across cell lines and tissues
**S3** - Compound number and class balance across ToxCast endpoint categories
**S4** - Assay PRAUC across representation and consensus profile strategy

***Figure S1.*** *Mean cell count (A), normalized LDH (B), and normalized MT (C) for each well position across a representative batch (prod_27).*

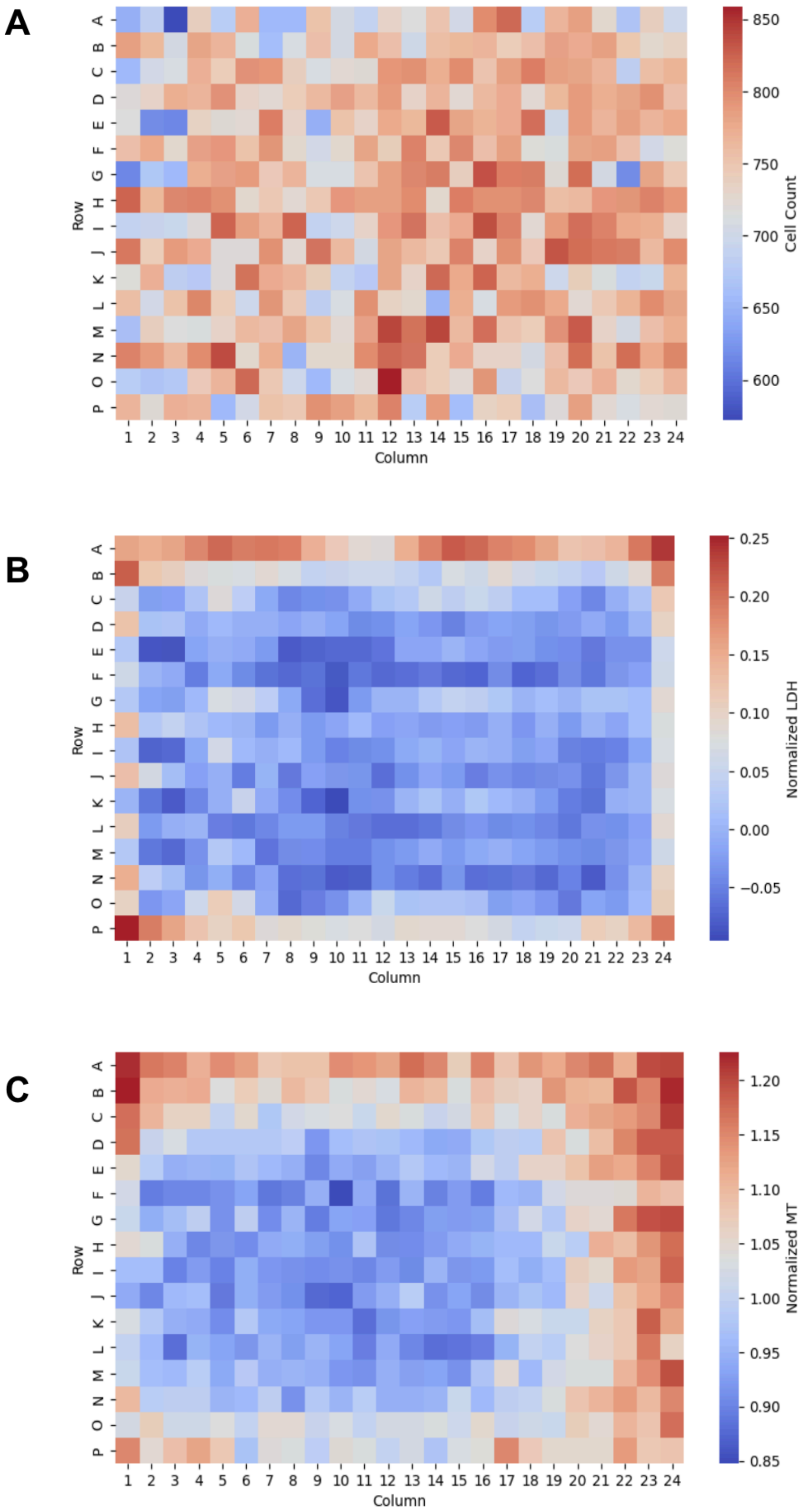

***Figure S2.*** *Cytotoxicity AC50 values across the 12 cell lines that had >800 OASIS compounds. Rows are compounds, columns are cell lines. Compounds that either had an AC50 > 100 or that did not have a detected AC50 were set to 100 (upper range of OASIS concentrations). Lower values (darker cells) indicate greater cytotoxicity in that it occurred at a lower concentration.*

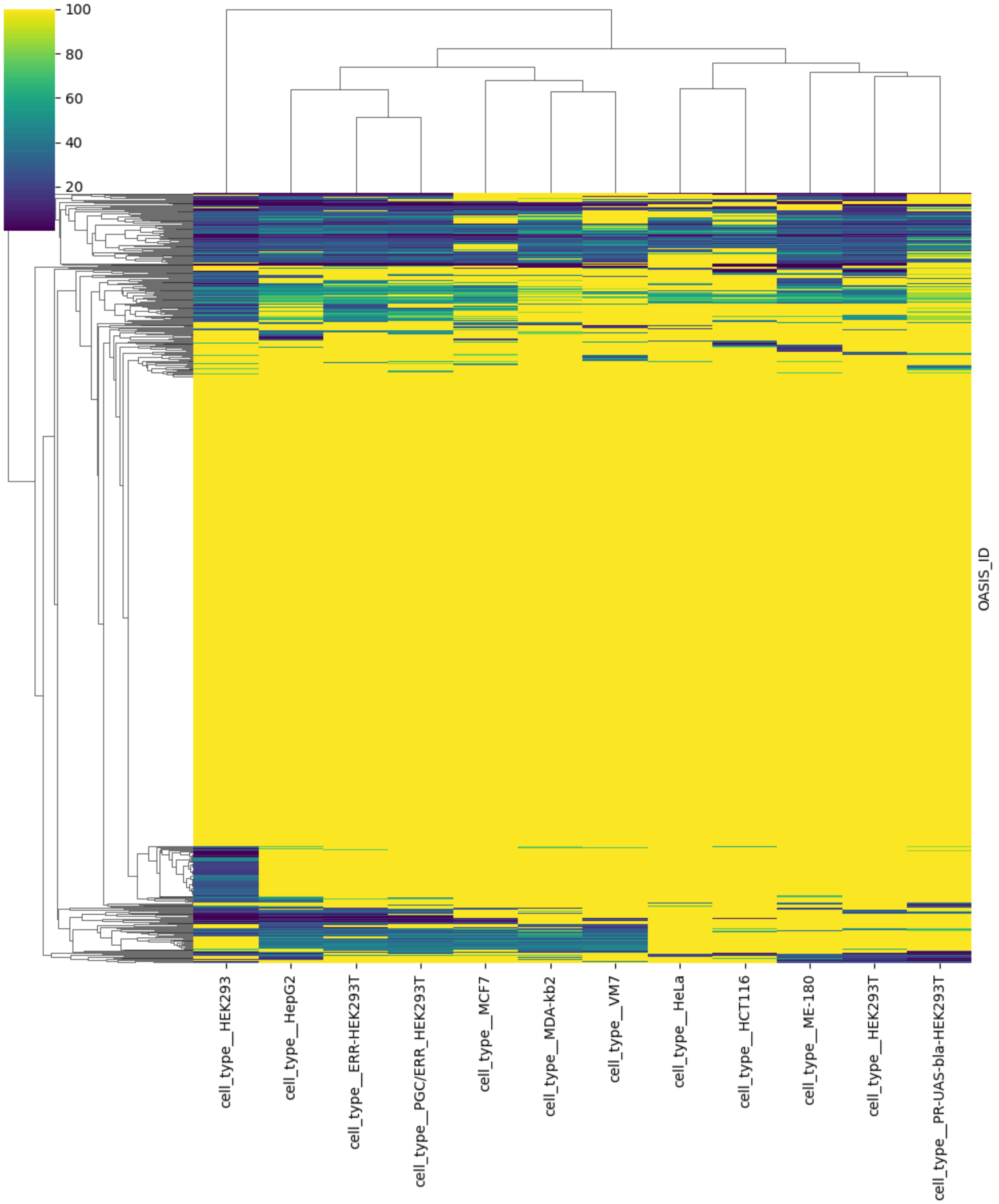

***Figure S3.*** *Distributions of the total number of tested compounds and the percentage of positive hits for endpoints from both the cytotoxicity, cell-based and cell-free assays.*

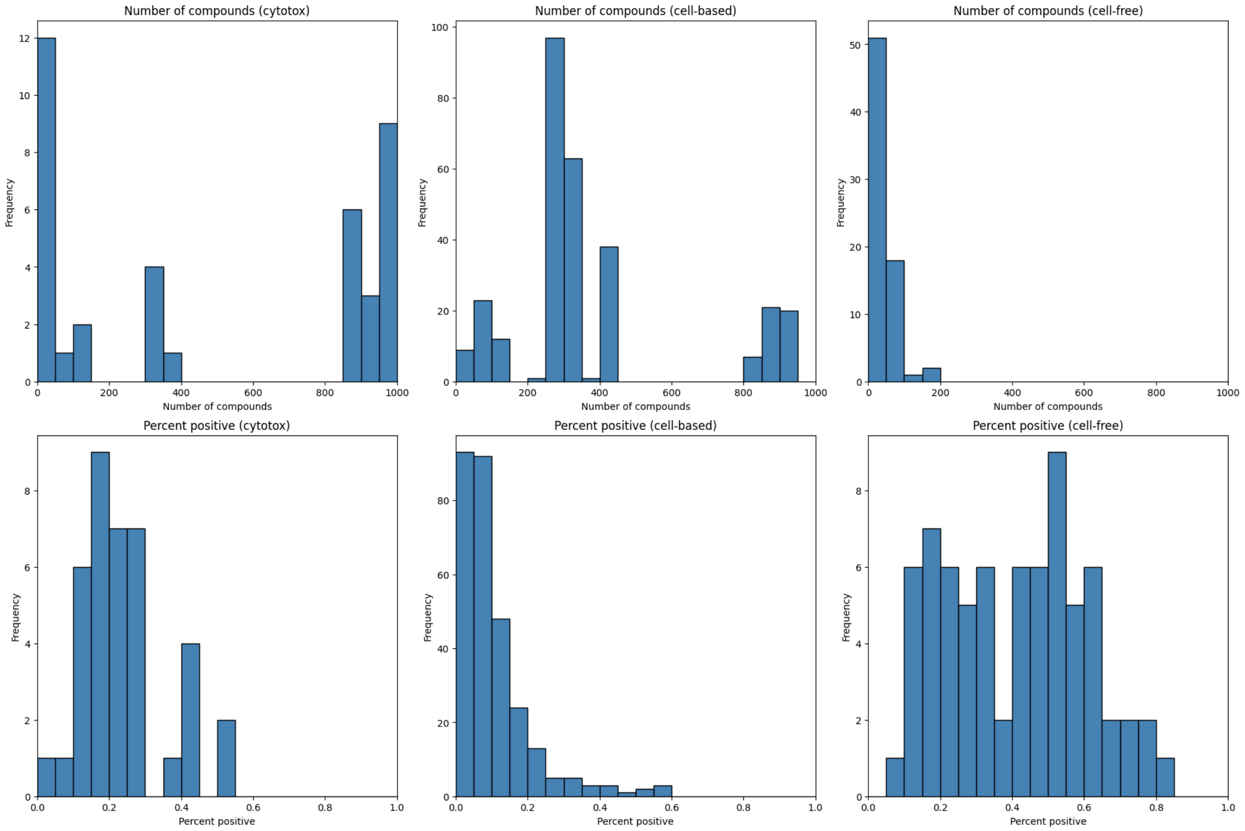

***Figure S4.*** *Classifier PRAUC, stratified by endpoint type, across consensus profile strategy (A), and cell representation (B).*

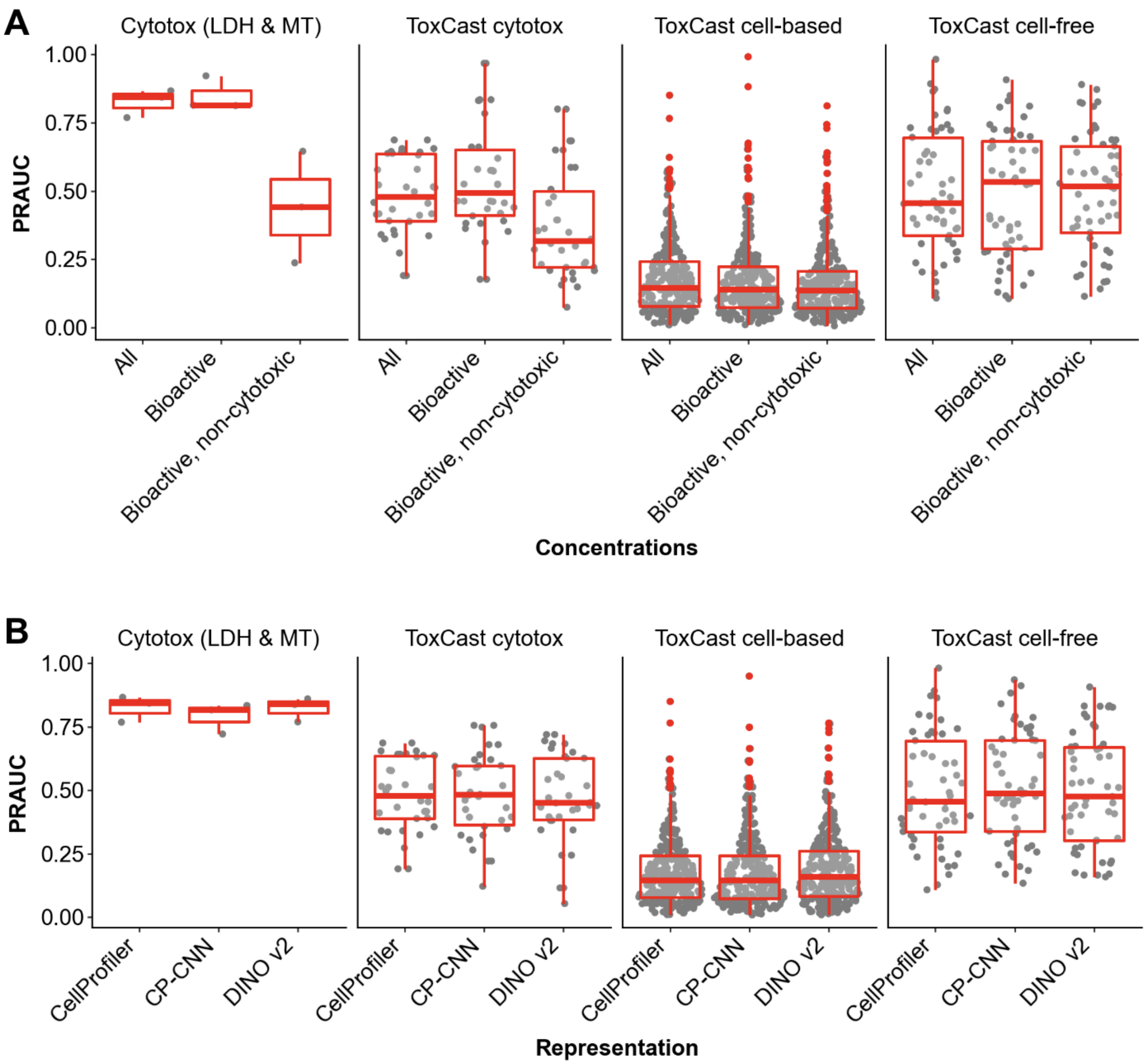
